# Unraveling factors responsible for pathogenic differences in Lassa virus strains

**DOI:** 10.1101/2024.05.21.595091

**Authors:** Satoshi Taniguchi, Takeshi Saito, Ruchi Paroha, Cheng Huang, Slobodan Paessler, Junki Maruyama

## Abstract

Lassa virus (LASV) is the etiological agent of Lassa fever (LF), a severe hemorrhagic disease with potential for lethal outcomes. Apart from acute symptoms, LF survivors often endure long-term complications, notably hearing loss, which significantly impacts their quality of life and socioeconomic status in endemic regions of West Africa. Classified as a Risk Group 4 agent, LASV poses a substantial public health threat in affected areas. Our laboratory previously developed a novel lethal guinea pig model of LF utilizing the clinical isolate LASV strain LF2384. However, the specific pathogenic factors underlying LF2384 infection in guinea pigs remained elusive. In this study, we aimed to elucidate the differences in the immunological response induced by LF2384 and LF2350, another LASV isolate from a non-lethal LF case within the same outbreak. Through comprehensive immunological gene profiling, we compared the expression kinetics of key genes in guinea pigs infected with LASV LF2384 and LF2350. Our analysis revealed differential expression patterns for several immunological genes, including CD94, CD19-2, CD23, IL-7, and CIITA, during LF2384 and LF2350 infection. Moreover, through the generation of recombinant LASVs, we sought to identify the specific viral genes responsible for the observed pathogenic differences between LF2384 and LF2350. Our investigations pinpointed the L protein as a crucial determinant of pathogenicity in guinea pigs infected with LASV LF2384.

**Author summary:** Lassa virus (LASV) is known to cause lethal hemorrhagic fever, Lassa fever, and is classified as a Risk Group-4 agent, which need to be handled in the highest biocontainment laboratories, biosafety level-4 (BSL-4), due to its high pathogenicity and the lack of preventive or treatment methods. LASV infection has a huge impact on public health and socioeconomics in endemic areas, however, its pathogenic mechanism is still largely unknown. In order to unveil the mechanisms of LASV pathogenesis, we compared the pathogenicity of two LASV isolates, which have opposite phenotypes in guinea pigs. Additionally, we determined the viral factor responsible for pathogenic differences between LASV isolates using reverse genetics. In summary, our study provides valuable insights into the immunological and virological factors underlying the pathogenic differences between LASV strains associated with lethal and non-lethal LF cases. Understanding these distinctions is essential for informing strategies for the diagnosis, treatment, and prevention of LF.

## Introduction

Lassa virus (LASV) belongs to the order *Bunyavirales*, family *Arenaviride* and is the causative agent of Lassa fever (LF). LASV is endemic in West African countries such as Nigeria, Guinea, Liberia, and Sierra Leone, and outbreaks occur annually. LF is a major public health threat in endemic regions of West Africa and up to 500,000 LASV infections are reported annually [1]. The case-fatality rate for hospitalized cases ranges from 15-70% depending on the outbreak [2–4]. Although an estimated 37.7 million people are at risk for LF, countermeasures against the virus are extremely limited [5]. Due to its high pathogenicity as well as a lack of countermeasures, such as vaccines and therapeutics, LASV is classified as a Risk Group 4 agent, and must be handled in the highest biological containment facility, biosafety level-4 (BSL-4). Requirement of BSL-4 facilities to handle the LASV imposes significant logistical and resource challenges on research aimed to develop preventive or therapeutic measures for LF. In order to develop effective countermeasures against LASV, understanding the viral factors responsible for its pathogenicity is essential.

Animal models of LF are limited, especially models utilizing clinical isolates representing the various different lineages of LASV. The nonhuman primate model, which is the gold standard for LF studies, correlates well with lethal human LF [6,7]. However, there is a concern about the high cost and numerous safety issues when handling them in high-containment laboratories. Recently, our laboratory developed a novel guinea pig model of LF [8]. In this model, outbred Hartley guinea pigs develop a uniformly lethal infection when infected with a clinical isolate of LASV from a lethal LF case, strain LF2384, without any host-virus adaptation [8]. We also found that the LASV LF2350, which was isolated from a non-lethal LF case in the same outbreak, did not induce a lethal infection in guinea pigs. These results indicated that our novel guinea pig LF model could be used to study differences in virulence of various LASVs and it recapitulates human clinical data. This valuable tool will allow us to study the pathogenic factors of LASV using a combination of molecular biology and *in vivo* studies.

In this present study, we utilized this model to explore the host and virus factors that contribute to the different pathogenicity between LF2384 and LF2350. Our assessment of the pathogenicity of recombinant LASV generated using reverse genetics approaches revealed that LASV RNA-dependent RNA polymerase (L) is the determinant viral factor of pathogenic differences between LASV LF2384 and LF2350. The findings presented in this paper provide insights into the molecular mechanisms underlying LASV infection, thereby accelerating the development of preventive and therapeutic methods against LF.

## Results

### Pathogenicity of LASV LF2350 in guinea pigs

LASVs LF2384 and LF2350 were isolated from serum samples collected in 2012 Sierra Leone outbreak. In our previous report, LASV LF2350 showed lower pathogenicity than LF2384 in Stat1-KO mice [9]. We also reported that LASV LF2384 caused lethal infection in Hartley guinea pigs [8]. To compare pathogenicity of LASVs LF2384 and LF2350 in immunocompetent animals, we investigated the pathogenicity of LASV LF2350 in Hartley guinea pigs. The guinea pigs were infected with 10^5^ or 10^4^ plaque forming unit (PFU) of LASV LF2350. We have observed that the guinea pigs infected with either dose exhibited minimal weight loss and only experienced transient fever. Importantly, these animals ultimately survived the infection without exhibiting any signs of disease (Fig. 1). This outcome is significantly opposite compared to LASV strain LF2384, which has been associated with severe disease and uniform lethality in guinea pigs. Previously we have reported that LASV LF2384-infected guinea pigs typically showed pronounced disease symptoms and succumbed to the infection [8]. The contrast outcomes between LF2384 and LF2350 on the guinea pigs suggests that these two LASV isolates have distinctly different phenotypes in Hartley guinea pigs.

**Fig. 1.**
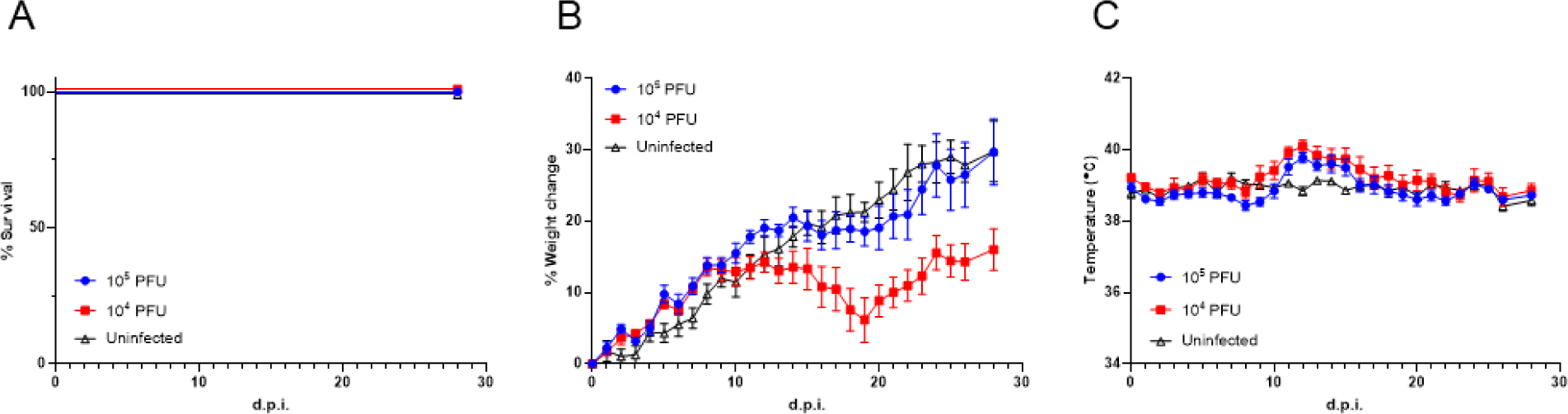
Pathogenicity of LASV LF2350 in Hartley guinea pigs. (A) Survival rate, (B) body weigh change, and (C) body temperature of Hartley guinea pigs infected with 10^4^ or 10^5^ PFU of LASV LF2350 (n=5) or uninfected were plotted, respectively. The error bars indicate standard errors.

### Hematology and virus dissemination of LASVs LF2384 and LF2350 in guinea pigs

Blood and organ samples were collected from guinea pigs infected with 10^4^ PFU of LASV LF2384 or LF2350 at 11 days post-infection (d.p.i.) to assess complete blood count (CBC), blood clinical chemistry, and virus dissemination (Fig. 2, Table S1, and S2) since guinea pigs infected with LASVs showed fever and weight loss around 11 d.p.i. (Fig. S1). Compared to the uninfected control group, guinea pigs infected with LF2384 or LF2350 exhibited leukopenia, lymphopenia, and thrombopenia. However, there was no significant difference between the samples infected with LASV LF2384 and LF2350. Likewise, the results of blood clinical chemistry and virus titers in organs or blood did not have any dramatical differences. Overall, these findings provide valuable insights into the pathophysiological consequences of LASV infection in guinea pigs and highlight the similarities between strains LF2384 and LF2350 in terms of their effects on host physiology and viral dissemination.

**Fig. 2.**
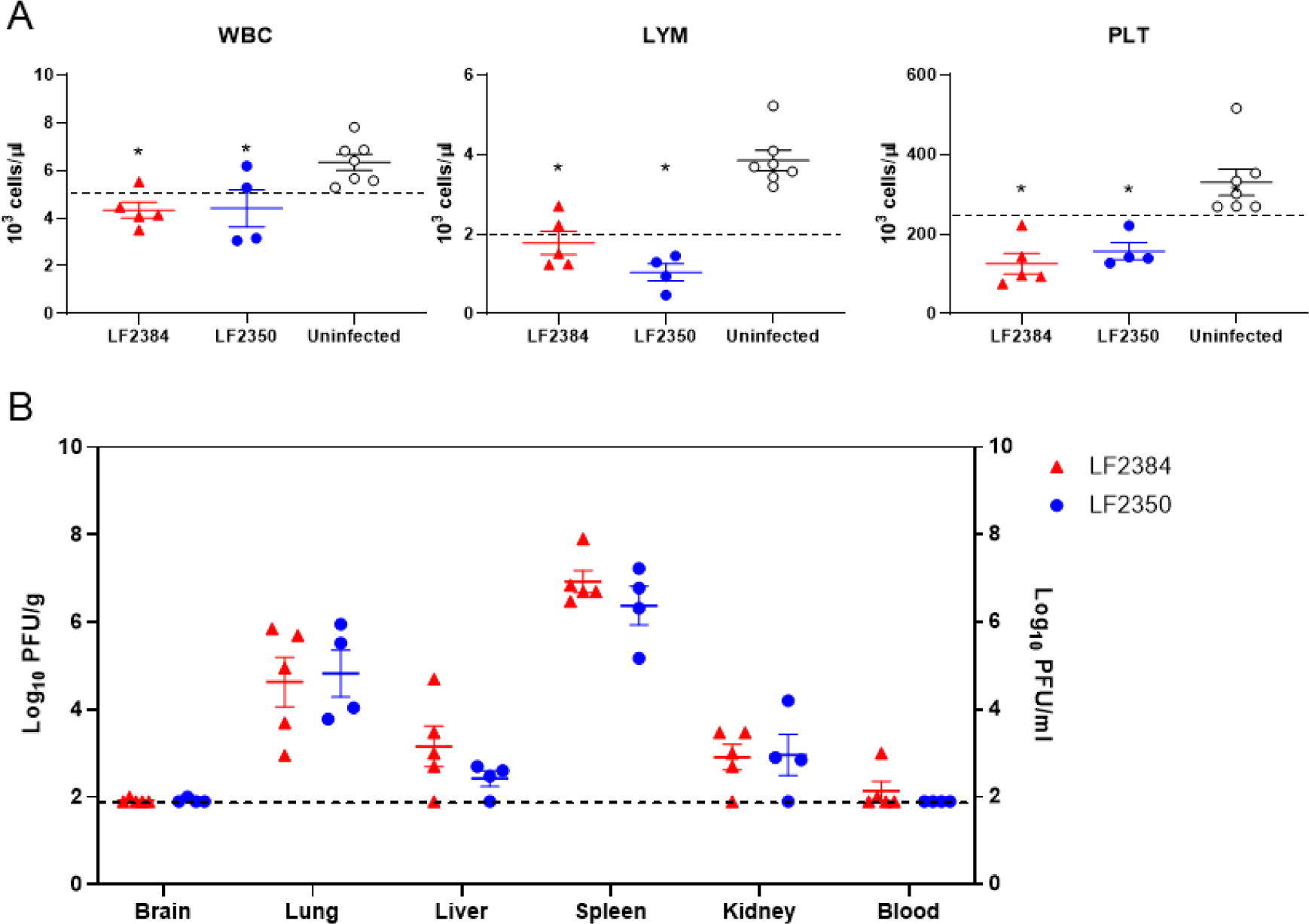
Complete blood counts and virus dissemination in guinea pigs infected with LASVs at 11 d.p.i. (A) White blood cells (WBC), lymphocyte (LYM), and platelet (PLT) were counted in blood samples from guinea pigs infected with 10^4^ PFU of LASV LF2384 or LF2350 at 11 d.p.i. or uninfected control. Each symbol represents the value for an individual guinea pig. The means ± standard errors for the groups are shown. The broken lines indicate the lower limit of the normal range. Statistical analyses were performed using One-way ANOVA following Šídák’s multiple comparisons test (p<0.05 compared to uninfected control). (B) Virus titers were measured in organ and blood samples collected from guinea pigs infected with 10^4^ PFU of LASV LF2384 or LF2350 at 11 d.p.i. Each symbol represents the value for an individual guinea pig. The means and standard errors were shown. The broken line indicates the detection limit (<2.0 log10 PFU/g for organs, <2.0 log10 PFU/ml for blood).

### Transcription profiling of immunological genes

Peripheral blood mononuclear cells (PBMCs) were collected from guinea pigs infected with 10^4^ PFU of LASV LF2384 or LF2350 at 11 d.p.i. and used to analyze transcription profiling of 93 genes related to immune system (Fig. 3 and Table S3). We found statistical differences (p<0.05) in 6 transcripts in PBMCs infected with LASV LF2384 and LF2350 (Fig. 3B). Notably, LASV LF2384 exhibited lesser downregulation of CD19-2 (B-lymphocyte antigen) compared to LASV LF2350. Conversely, LASV LF2384 downregulated CD94 (a natural killer cell marker), whereas LF2350 showed a slight upregulation of CD94. Additionally, LASV LF2384 exhibited lesser downregulation of CD23 (Fcε-receptor II) compared to LF2350. Furthermore, we have observed that CD92 (Choline transporter-like protein 1) was more prominently upregulated by LASV LF2384 infection compared to LF2350. Additionally, interleukin-7 (IL-7) was upregulated by LASV LF2384 infection in contrast to LF2350. The expression of CIITA, a crucial gene involved in regulating major histocompatibility complex (MHC) class II expression, displayed intriguing differences between the two LASV strains. CIITA was upregulated by LASV LF2384 infection, whereas LF2350 infection led to its downregulation. These findings underscore the remarkable differences in the host immune response elicited by LASV strains LF2384 and LF2350. Which provide valuable insight for understanding the molecular mechanisms underlying LASV infection and the subsequent host immune response.

**Fig. 3.**
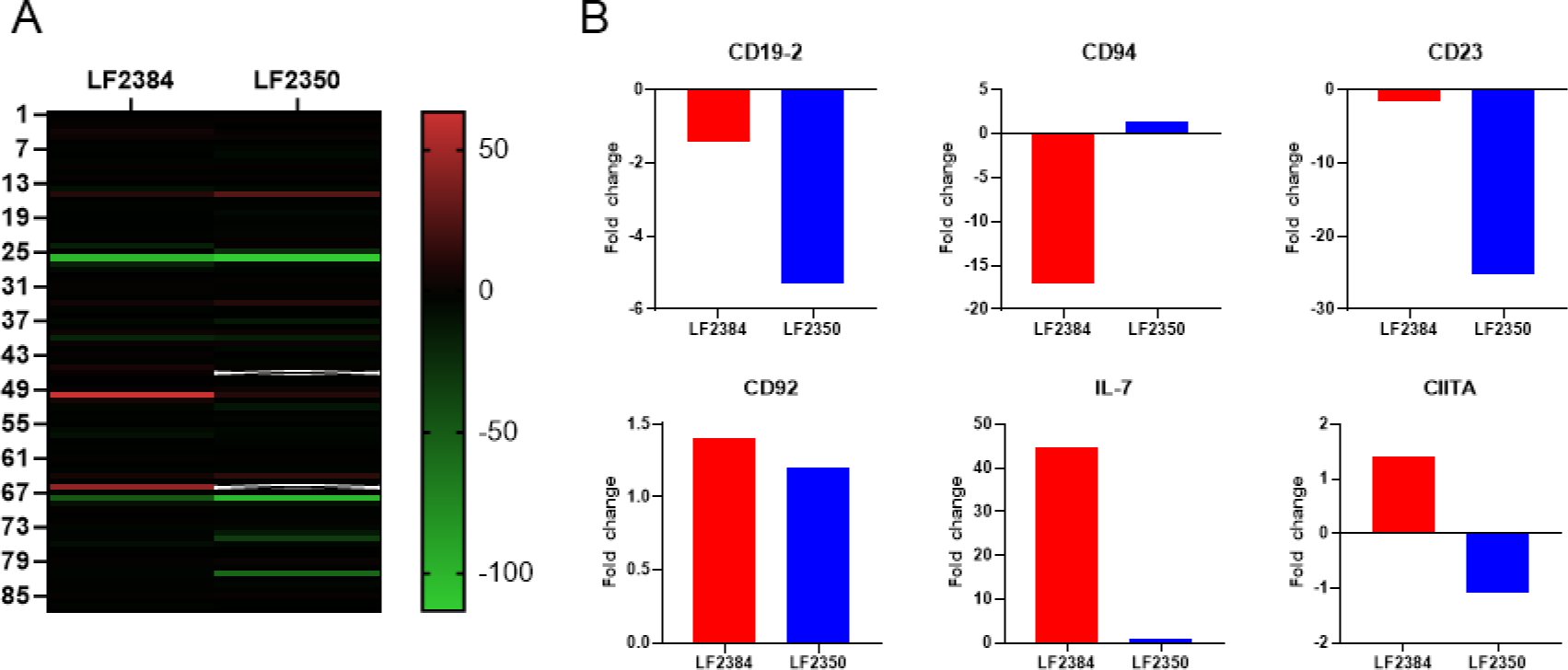
Transcription profiling of immunological genes in guinea pigs infected LASVs. (A) Transcription profiling of Heatmap of 94 genes related to immune response was presented as a heatmap. (B) Transcription changes of genes that have statistical differences between guinea pigs infected with LASV LF2384 and LF2350 at 11 d.p.i. (p<0.05) using delta-delta Ct method. Fold changes in guinea pigs infected with 10^4^ PFU of LASV LF2384 or LF2350 at 11 d.p.i. were compared to uninfected control.

### The viral factor responsible for pathogenic differences between LASV LF2384 and LF2350

To determine the viral factor responsible for LASV pathogenicity in guinea pigs, we rescued recombinant LASVs (rLASVs) by using reverse genetics system and evaluated their pathogenicity in guinea pigs. Generated rLASVs are summarized in Fig. 4A. Recombinant LASV LF2384 (r2384) and LASV LF2350 (r2350) had similar pathogenicity compared to the parent wildtype viruses (Fig. 4B). At the same time, we assessed the pathogenicity of the recombinant segment-swapped viruses, rS50L84 and rS84L50. rS50L84, which has S-segment of LASV LF2350 and L-segment of LASV LF2384, showed similar pathogenicity in guinea pigs compared to wildtype LASV LF2384 or r2384, indicating that the viral protein encoded in L-segment, Z and/or L is responsible for pathogenic differences of the LASVs in guinea pigs (Fig. 4B). Next, we generated rLASVs which open-reading frame of L was swapped, r2350L84 and r2384L50, and evaluated their pathogenicity in guinea pigs (Fig. 4C). r2384L50 did not show pathogenicity similar to r2350, whereas r2350L84 showed 100% lethality. These results indicated that LASV L protein is a pathogenic/attenuation factor in guinea pigs.

**Fig. 4.**
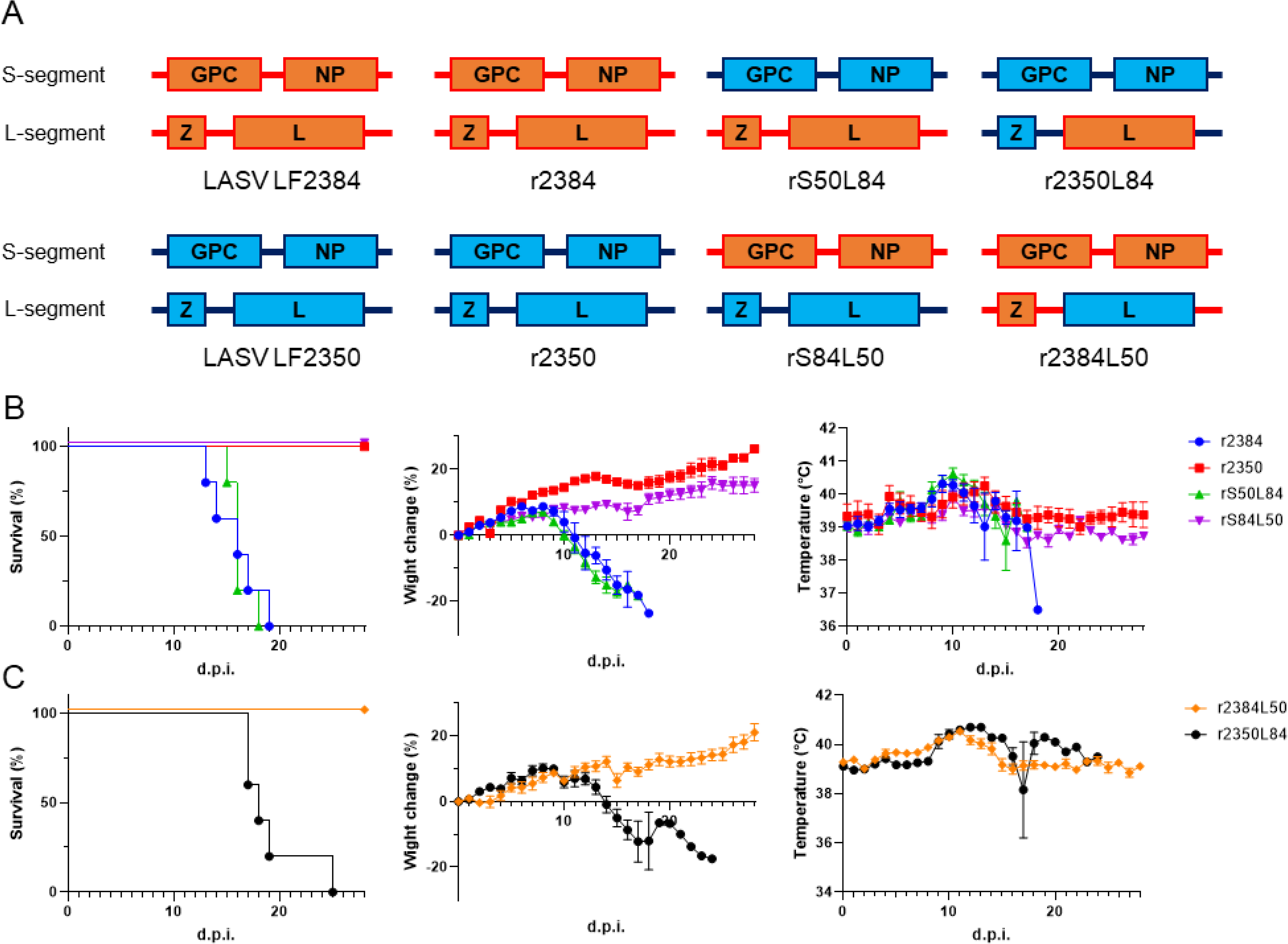
Pathogenicity of rLASVs in guinea pigs. (A) Schematic figures of recombinant LASVs. (B) Survival rate, body weigh change, and body temperature of Hartley guinea pigs infected with 10^4^ PFU of r2384, r2350, rS50L84, or rS84L50 (n=5). Means and standard errors were plotted. (C) Survival rate, body weigh change, and body temperature of Hartley guinea pigs infected with 10^4^ PFU of r2384L50 or r2350L84 (n=5). Means and standard errors were presented.

**Fig. 5.**
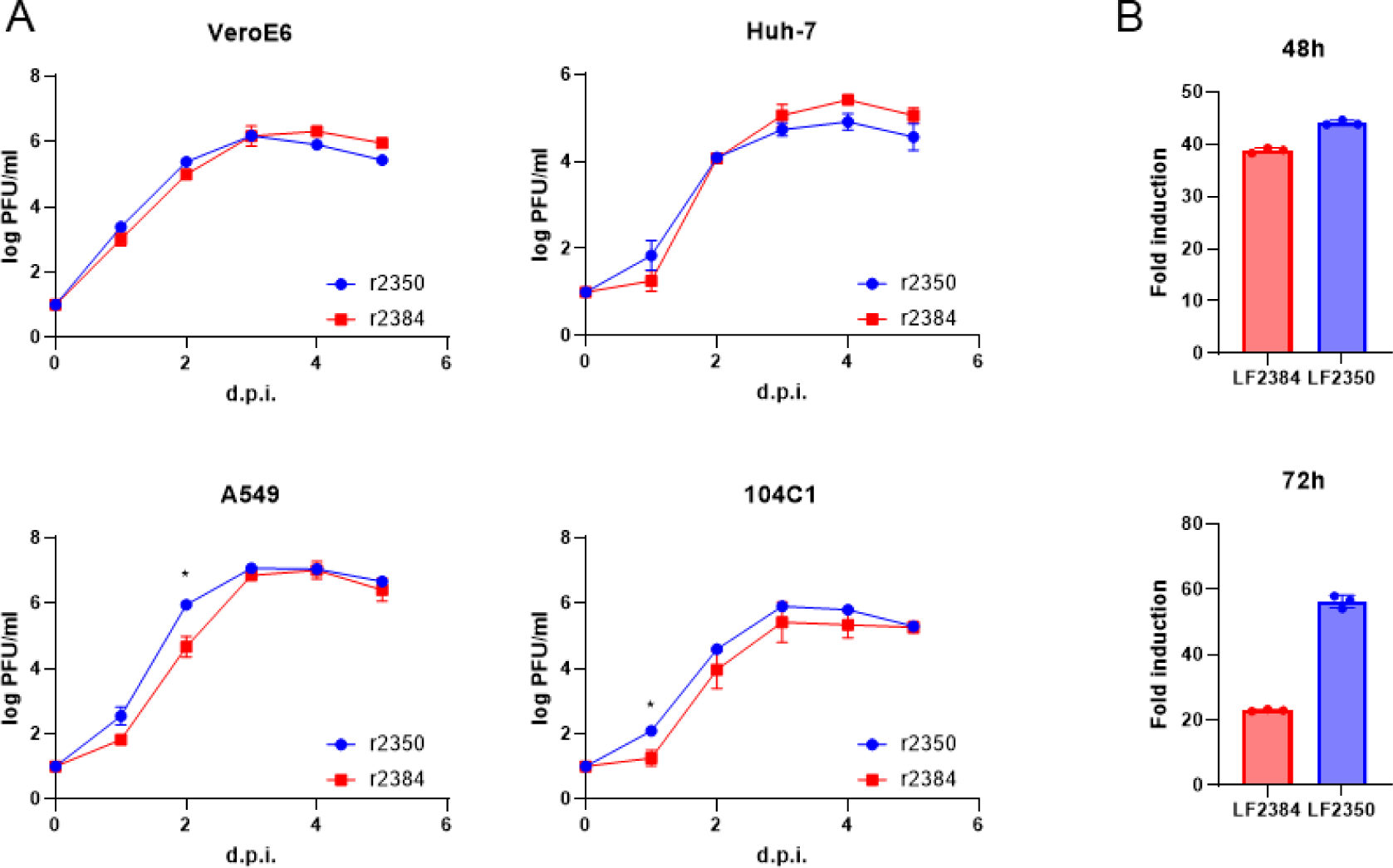
Virus growth kinetics and RdRp function of LASV LF2384 and LF2350. (A) The virus growth kinetics of r2384 and r2350 was measured in Vero E6, Huh-7, A549, and 104C1 cells. The graphs were shown in means and standard deviations of biological triplicates. (B) The minigenome reporter assay was performed to compare RdRp functions of LASV LF2384 and LF2350. The fold change in luciferase activity was calculated by luminescence in cells transfected with the same amount of minigenome plasmid and empty pCAGGS vector as a standard.

### Virus replication kinetics and RdRp function of LASV LF2384 and LF2350

Due to its role in differences of virus pathogenicity between LASV LF2384 and LF2350 in guinea pigs, we investigated the RdRp function of LASV LF2384 and LF2350 and their virus replication kinetics in different cell lines. The virus growth kinetics of r2384 and r2350 in non-human primate (Vero E6), human (Huh-7 and A549), and guinea pig (104C1) cell lines do not have any dramatical differences (Fig. 6A). These results are consistent with virus dissemination data *in vivo* (Fig. 2B). However, the results of minigenome reporter assay revealed that LASV LF2350 had higher RdRp function than LASV LF2384 (Fig. 6B). This data suggests that despite the similar virus growth kinetics observed *in vitro* and *in vivo*, there are clear differences in the RdRp activity between the two LASV strains.

## Discussion

LF is one of the most severe public health threats in both endemic regions of West Africa and worldwide due to its high morbidity and mortality rates [2,10]. There have been several imported cases of LF to the United States and Europe via infected travelers [11,12]. In addition, one-third of Lassa fever survivors develop sensorineural hearing loss, which is often permanent, leading to a huge impact on economic and the quality of life for survivors [13]. Although LASV is one of the most alarming pathogens from a public health perspective, there are no licensed vaccines and few therapeutics against LF. Basic scientific research is essential to aid the development of effective countermeasures against infectious diseases. However, most studies of LASV and LF are primarily focused on either countermeasure development or in vitro molecular biology, as live LASV must be handled in BSL-4 facilities. In this manuscript, we revealed the host and viral factor responsible for LASV pathogenesis using recently developed guinea pig models of LF [8].

LASVs are divided into seven lineages based on their phylogenetic relationships [14]. In this study, we compared the pathogenicity of two LASV isolates, LF2384 and LF2350, which belong to lineage IV, in guinea pigs. LASV LF2384 was isolated from a lethal LF case, and LF2350 was isolated from a non-lethal case [9,15]. Interestingly, the pathogenicity of LASVs in guinea pigs linked to the lethal outcome in human patients. The pathogenicity of these LASVs in Stat1-KO mice has the similar trend [9,15].

There are no dramatical differences in CBC, blood clinical chemistry, and virus dissemination of guinea pigs infected with 10^4^ PFU of LASV LF2384 and LF2350 at 11 d.p.i. although disease outcomes of these viruses are completely opposite. Transcription analysis of genes related to the immune response in PBMCs revealed that several differences. CD94 was downregulated in guinea pigs infected with LASV LF2384, whereas it was slightly upregulated in guinea pigs infected with LF2350 (22.8-fold differences between LF2384 vs LF2350). CD94 is expressed in NK cells and a subset of CD8+ T-cells [16] and is essential for NK cell mediated resistance against viral infection [17]. Our results indicated that decrease or less activation of NK cells was responsible for lethal outcomes of LASV infections. This hypothesis is consistent with previous reports that cellular immunity-related virus clearance is important for protection against LASV infection [18–20]. It was reported that Lujo virus, another member of arenaviruses causing hemorrhagic fever in African continent, caused nonlethal infections in macaques, possibly due to early mobilization of NK cells might be involved in reduced pathogenicity [21]. In LASV LF2350-infected guinea pigs, NK cells might be also mobilized in the early stage of the infection, resulting in the low pathogenicity. Furthermore downregulation of CD19-2 (B-cell antigen) and CD23 (Fcε-receptor II) was observed in LASV LF2350-infected animals than LASV LF2384, indicating that more B-cell decrease in LASV LF2350-infected animals. IL-7 upregulation was observed only in guinea pigs infected with LASV LF2384. IL-7 can restore B cell numbers [22]. These results indicated that B-cells decreased in guinea pigs infected with both LASVs, however B-cells were restored via IL-7 production only in guinea pigs infected with LASV LF2384. CIITA (MHC class II transactivator) regulates major histocompatibility complex (MHC) class II expression in antigen presenting cells (APCs) [23] It was reported that CIITA could play preventive or supportive roles in virus infections [24]. Our data indicated that CIITA activation led to a lethal outcome in LASV infection, suggesting a complex interplay between host immune responses and virus pathogenesis. Overall, these findings highlight the intricate dynamics of the host immune response in LASV infection and its implications for disease pathogenesis. Therefore, further studies are desirable to unveil host immune response in LASV infection related to its pathogenesis.

We also identified the L protein as a key determinant of pathogenic differences between LASV strains LF2384 and LF2350, using reverse genetics and in vivo experimentation. As the largest protein in arenaviruses, the L protein functions as an RNA-dependent RNA polymerase (RdRp), governing RNA synthesis and transcription in negative-strand RNA viruses. Surprisingly, despite LF2350 exhibiting superior RdRp function in minigenome reporter assays, virus growth kinetics did not differ in various cell lines derived from human, non-human primate, or guinea pig. These findings suggest the presence of unidentified functions in the L protein that may be relevant to pathogenicity.

Several studies reported L segment was virulent factor for arenaviruses such as Pichinde virus (PICV), Lymphocytic choriomeningitis virus, and ML-29 [25–27]. Especially, the study conducted by McLay L et al. suggested that substitutions of 4 amino acids located in C-terminal domain of L (T1808A, L1839V, N1889D, and N1906D) attenuated the virulence of PICV *in vivo*. The study revealed that these substitutions decreased replicative efficiency of RdRp, resulting in the attenuation of virulence *in vivo*. Interestingly, the replicative efficiency of LF2384 was lower than that of LF2350, suggesting that the relation between virulence *in vivo* and replicative efficiency were completely opposite to the results of that PICV research. In this respect, the pathogenic factor of LASV LF2384 L protein is completely different from that of PICV L protein reported by McLay L et al.

Recently crystal structure of LASV L was reported and putative function domains were determined [28]. L protein consists of 2220 amino acids. One hundred twenty-one amino acid differences were found in LF2384 and LF2350 L proteins (Table 1). Amino acid differences were found in all putative domains and the amino acid identity rate in these domains between LASV LF2384 and LF235 were above 90%, suggesting that all L protein open reading frame regions have the possibility of responsible region of virulence. Despite a high amino acid identity rate in putative domains, indicating the possibility of multiple regions contributing to virulence, the rescued recombinant LASVs with chimeric L proteins did not confer pathogenicity in guinea pigs (data not shown). This suggests a complex interplay between the L protein and host factors in determining LASV pathogenicity.

**Table 1.** Amino acid differences in L between LASV LF2384 and LF2350.

|  | Domain |  |  |  |  |  |  |
| --- | --- | --- | --- | --- | --- | --- | --- |
|  | Endonuclease | Linker | PA-C-like region | RdRp | PB2-N-like region | C terminal domain | Total |
| Number of variant amino acid | 5 | 2 | 40 | 34 | 4 | 36 | 121 |
| Putative number of amino acid | 194 | 66 | 450 | 902 | 209 | 399 | 2220 |
| Identity (%) | 97.4 | 97.0 | 91.1 | 96.2 | 98.1 | 91.0 | 94.5 |

Given the transcription analysis, the differences in L are expected to contribute to the differences in regulation of CD19-2, CD94, CD23, CD92, IL-7, and CIITA, but more detailed analysis is needed to determine the mechanism of different regulation of each factor and how they contribute to LASV pathogenicity. Notably, previous studies on the prototypic LASV strain Josiah in guinea pigs suggested that the S segment, rather than the L segment, serves as a main virulent factor. [29]. In that study, one of the main virulent factors of LASV Josiah strain, a prototypic strain of LASV, in guinea pig infection model was concluded to not L segment but S segment. However, differences in strain, host model, and infection route between our study and prior research may account for disparate conclusions. Further investigations are warranted to elucidate the direct or indirect roles of the L protein of LF2384 or the S segment of Josiah in interacting with host factors in guinea pigs.

In summary, our findings underscore the multifaceted nature of LASV pathogenicity and highlight the need for continued research to unravel the intricate mechanisms governing virulence in Lassa fever.

## Materials and methods

### Cell and Virus

Vero, Vero E6, A549, Huh-7, and 104C1 cells were maintained in Dulbecco’s modified Eagle’s medium (DMEM) supplemented with 10% fetal bovine serum (FBS) and 100 U/mL of penicillin-streptomycin. BHK-21S cells were maintained in DMEM supplemented with 5% FBS, 10% tryptose phosphate broth (Gibco), 100 U/mL of penicillin–streptomycin, and 2 mM of L-glutamine. All cell lines were cultured at 37°C in 5% CO_2_.

LASV LF2384 (Gene accession number: pending) and LF2350 (Gene accession number: pending), which belong to lineage IV, were isolated from serum samples of human LF cases during a 2012 outbreak in Sierra Leone [8,9]. The viruses were propagated in Vero cells, and virus-containing cell culture supernatant was stored in -80°C freezer until use. All work with infectious LASV was performed in biosafety level 4 (BSL-4) facilities in the Galveston National Laboratory (GNL) at the University of Texas Medical Branch (UTMB) in accordance with institutional guidelines.

### Virus titration

Virus titers were determined by plaque assay as described elsewhere [8]. Briefly, confluent monolayers of Vero cells in 12-well plates were inoculated with 100 µl of 10-fold diluted virus and incubated for 30 minutes at 37°C in CO_2_ incubator. After removing the inoculum, each well was overlaid with minimum essential medium containing 2% FBS, 1% penicillin-streptomycin, and 0.6% tragacanth (Sigma). Following a 5- to 6-day incubation, cells were fixed with 10% Formalin and stained with Crystal Violet. Viral titers were represented as plaque forming unit (PFU).

### Virus challenge into guinea pigs

Six-to eight-week-old Hartley guinea pigs were purchased from Charls River Laboratories. All animals were housed in ABSL-2 and ABSL-4 facilities in the GNL at UTMB. All animal studies were reviewed and approved by the Institutional Animal Care and Use Committee at UTMB and were carried out according to the National Institutes of Health guidelines. Animal identification and measuring of body temperature were performed with subcutaneously implanted BMDS IPTT-300 transponders and a DAS-8027 transponder reader (Bio Medic Data Systems). Guinea pigs were intraperitoneally inoculated with diluted viruses in 100 µl of PBS or 100 µl of PBS as an uninfected control. Inoculation titers of recombinant LASVs in animal infection experiments were unified to 10^4^ PFU/ head. Guinea pigs were monitored illness with measuring body weight and temperature.

### Complete blood count and blood clinical chemistry

Whole-blood samples were collected in EDTA tubes from guinea pigs infected with LASV LF2384 or LF2350 at 11 d.p.i. Complete blood counts were measured using a VETSCAN HM5 (Zoetis) according to the manufacturer’s instructions. Blood clinical chemistry was measured with plasma samples using VETSCAN VS2 and Comprehensive Diagnostic Profile (Zoetis) according to the manufacturer’s instructions.

### Immunological PCR array

Transcription of immunological gene was analyzed by PCR assay according to previously described [30]. Briefly, Peripheral blood mononuclear cells (PBMCs) were isolated from blood samples from guinea pigs infected with LASV LF2384 or LF2350 at 11 d.p.i. by using Ficoll-paque Premium 1.084 (Sigma) according to manufacturer’s instruction. RNA was extracted from PBMCs using TRIzol reagent (Thermo Fisher Scientific) and Roche cellular RNA large volume kit in the MagNA Pure 96 instrument. cDNA was produced using the BioRad iScript system according to the manufacturer’s instructions. The immunological PCR array was performed as previously described [31,32]. Ct values were analyzed using delta-delta Ct.

### Viral genome sequencing

Viral genome sequences of LASVs were determined by using Sanger sequencing. Total RNA was extracted from infected Vero cells. The open-reading flame of each viral protein and intergenic region was amplified by Reverse transcription polymerase chain reaction (RT-PCR). To determine the 5’ and 3’ termini of the viral genome sequence, a 5’ RACE system (Thermo Fisher Scientific, Carlsbad, CA) was used according to the previous study [33]. All amplified DNA fragments were purified, and their nucleic sequences were analyzed by Sanger sequencing.

### Plasmids

The S-segment and L-segment were cloned into pRF vector, which is controlled by murine RNA polymerase I (mPol-I) promoter and terminator to synthesis viral RNA. An additional G residue, which has been reported to enhance the efficiency of virus rescue by reverse genetics, was inserted between the mPol-I promoter and the viral genome sequences [34]. pRF vectors with chimeric L-segments were constructed by in-fusion high-fidelity (HD) restriction-free cloning method (TaKaRa Bio). Two nucleotides of LF2384 L gene were different from those of parent wildtype virus and one nucleotide of LF2350 GPC gene of was different from that of wildtype one. However, all of them were synonymous changes in amino acid sequence. The plasmids expressing LASV NP and L were constructed by cloning PCR amplicons of the LASV NP or LASV L genes into pCAGGS [35] linearized by EcoRI/XhoI or EcoRI/NheI using an in-fusion HD restriction-free cloning method. For minigenome plasmid, the NP gene was substituted for a Firefly luciferase (Fluc) gene, and the GPC gene was removed based on the pRF vector with LASV S-segment.

### Rescue of recombinant LASVs

Rescue of recombinant LASVs was performed according to the previous study [33]. Briefly, BHK-21S cells were seeded into wells of 12-well plate 1 day before transfection. Cells were transfected using 6.0 µL of FuGENE HD transfection reagent (Promega) with 0.4 µg of pRF-LASV-Sseg, 0.6 µg pRF-LASV-Lseg, 0.4 µg of pC-LASV-NP, and 0.6 µg of pC-LASV-L. Cells were incubated for 4-5 days. The medium used for culturing transfected BHK-21S cells was harvested and stored. Vero cells were inoculated with the medium for 5–6 days, and the supernatants were harvested and stored at -80°C. After viral propagation was confirmed using plaque assay, the RNA of each recombinant virus was extracted from infected cells, and the viral genome sequences of all recombinant viruses were confirmed using Sanger sequencing as described above.

### Comparison of virus growth

The virus growth kinetics was evaluated as previously described [33]. Briefly, Vero E6, Huh-7, A549, or 104C1 cells were infected with each virus at multiplicity of infection (MOI)=0.001. The cells were washed three times after a 30-min adsorption period at 37°C, and 1 mL of DMEM+2% FBS was added to each well. The cell culture supernatant samples were collected at 0–5 d.p.i. The collected supernatants were centrifuged at 8,000 × g for 5 min to remove cell debris and were stored at -80°C until the virus titer was measured by plaque assay.

### Minigenome reporter assay

The minigenome reporter assay was performed using a modified version of the protocol described previously [36]. Briefly, BHK-21S cells in 12-well plate were transfected with 0.25 µg of the minigenome plasmid, 0.25 µg of NP expression vector or pCAGGS vector, and 0.25 µg of L expression vector by using FuGENE HD transfection reagent (Promega). After the incubation for 48- or 72-hours post-transfection, firefly luciferase (Fluc) activities were measured using Luciferase Assay System (Promega). The luminescent was measured a GloMax 96 luminometer (Promega). The fold change in luciferase activity was calculated by luminescence in cells transfected with the same amount of minigenome plasmid and empty pCAGGS as a standard.

### Statistical analysis

Statistical analyses were performed with GraphPad Prism Software (ver. 9.1.2). Statistical differences were calculated by 2-way ANOVA followed by Dunnett’s post hoc test for body weight/temperature results. All viral growth curves were analyzed using a two-way ANOVA and Šidák’s multiple-comparison test to analyze the common logarithm of viral titers.

## Acknowledgment

The authors thank all the staff of the Animal Resources Center (ARC) at the University of Texas Medical Branch.

## Funding

National Institute of Health K99AI156012, R00AI156012 (JM)

National Institute of Health U01AI151801 (SP)

National Institute of Health R21AI166985 (CH)

The Institute for Human Infections and Immunity (JM)

John S Dunn Foundation (SP)

## Author contributions

Conceptualization: JM

Funding Acquisition: JM, SP

Methodology: JM, ST, TS

Investigation: JM, ST, TS, RP, JTM

Supervision: JM, SP

Writing – original draft: JM, ST, TS

Writing – review & editing: JM, ST, TS, RP, CH, SP

## Supporting information

**Table S1. Complete blood count in guinea pigs infected with LASVs at 11 d.p.i.**

**Table S2. Blood clinical chemistry in guinea pigs infected with LASVs at 11 d.p.i.**

**Table S3. Immunological transcription analysis in PBMCs**

**Fig. S1 Body weight and temperature change in guinea pigs infected with 10^4^ PFU of LASV LF2384 or LF2350.**

## Data and materials availability

All data are available in the main text or the supplementary materials. Sequences have been deposited in GenBank under accession no. pending.

## Supplementary

**Fig. S1.**
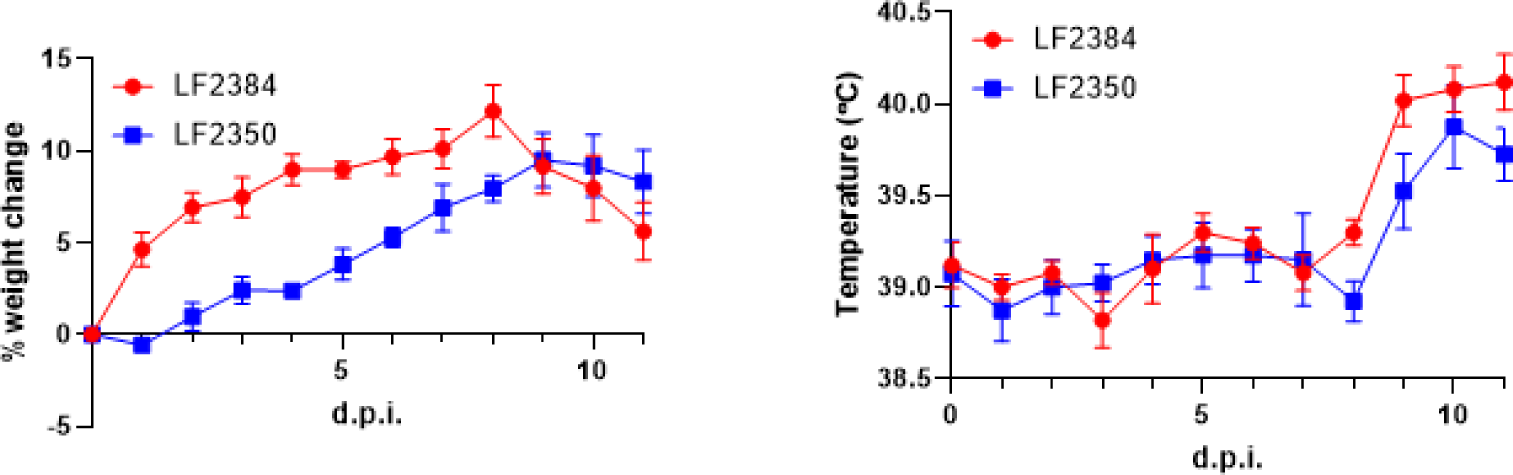
Body weight and temperature change in guinea pigs infected with 10^4^ PFU of LASV LF2384 or LF2350. Hartley guinea pigs were inoculated with 10^4^ PFU of LASV LF2384 or LF2350 intraperitoneally. Body weight (left) and temperature (right) were measured daily until 11 d.p.i. The means and standard errors were plotted. The number of animals were n=5 for LASV LF2384 and n=4 for LASV LF2350 since one guinea pig died after virus inoculation and blood sampling. Organ and blood samples was collected for CBC, virus dissemination, and transcription analyses.

